# Single-cell transcriptome analysis of the early immune response in the lymph nodes of *Borrelia burgdorferi*-infected mice

**DOI:** 10.1101/2023.11.29.569164

**Authors:** Varpu Rinne, Kirsi Gröndahl-Yli-Hannuksela, Ruth Fair-Mäkelä, Marko Salmi, Pia Rantakari, Tapio Lönnberg, Jukka Alinikula, Annukka Pietikäinen, Jukka Hytönen

## Abstract

Lyme borreliosis is a disease caused by *Borrelia burgdorferi* sensu lato bacteria. *Borrelia burgdorferi* is known to induce prolonged extrafollicular immune responses and abnormal germinal center formation. However, the mechanism behind this is poorly understood. The extrafollicular response is characterized by strong plasmablast induction and by an IgM, IgG3, and IgG2b dominant antibody production. These antibodies do not generate a neutralizing type of immunity, and the bacteria eventually establish a persistent infection. Here, we performed single-cell RNA sequencing to characterize the immune landscape of lymph node lymphocytes in the early *Borrelia burgdorferi* infection in a murine model.

Our results indicate that four days after *Borrelia burgdorferi* infection, a notable B cell proliferation, immunoglobulin class switching to IgG3 and IgG2b isotypes, and plasma cell differentiation are induced, all of which are hallmarks of the extrafollicular immune response. In addition, we found infection-derived upregulation of suppressor of cytokine signalling genes *Socs1* and *Socs3,* and downregulation of genes involved in MHC II antigen presentation in B cells.

Our results support the central role of B cells in the immune response of a *Borrelia burgdorferi* infection, and provide cues of mechanisms behind the determination between extrafollicular and germinal center responses during *Borrelia burgdorferi* infection.

## 1. Introduction

Lyme borreliosis, the most common vector-borne disease in Europe and the United States, is caused by *Borrelia burgdorferi* sensu lato bacteria (later: *B. burgdorferi*) [1,2]. A characteristic immune response to *B. burgdorferi* in its natural hosts serves to reduce and control the spirochete numbers and minimize the manifestations of the infection. However, the bacteria establish a persistent infection, enabling the re-acquisition of the spirochetes by the tick and their subsequent transmission to new hosts. Therefore, it seems likely that the co-evolution of *B. burgdorferi* and its reservoir host species has resulted in a state that maintains the enzootic cycle while helping the host species avoid overt disease manifestations [3,4]. On the other hand, some host species develop a symptomatic inflammatory response, which is seen also in humans. Indeed, without antibiotic treatment, Lyme borreliosis can persist also in humans for years. Whether this is due to a weak initial immune response or ineffective clearance of bacteria is not properly understood [5].

Early *B. burgdorferi* dissemination activates both innate and adaptive immune responses, resulting in macrophage and dendritic cell-mediated phagocytosis as well as eventually antibody-induced killing of the bacterial cells [3]. Immune response to *B. burgdorferi* infection is characterized by a lack of robust T-dependent B cell responses and impaired germinal center formation [5]. In contrast, *B. burgdorferi* induces a prolonged extrafollicular response by poorly understood mechanisms [6]. It has been suggested that extrafollicular immune response is driven by strong antigen recognition [7]. Indeed, in *B. burgdorferi* infection, intact spirochetes are widely present in lymph nodes already in the early phase of the infection [8,9]. The extrafollicular response to *B. burgdorferi* is characterized by strong plasmablast (PB) responses in lymph nodes, and especially by an IgM, IgG3, and IgG2b dominant antibody production [4,9]. However, these extrafollicularly derived antibodies do not generate a long-term neutralizing type of immunity [10,11].

Here, we have focused on the early-stage infection and have analysed single-cell RNA sequencing (scRNA-seq) profiles of draining lymph node lymphocytes from *B. burgdorferi* - infected mice. We provide evidence for plausible reduction of T cell – B cell interaction and bias for an extrafollicular response. The results give insight into the mechanisms behind the mainly extrafollicular *B. burgdorferi*-specific B cell response.

## 2. Materials and methods

### 2.1. Cell isolation, library preparation and sequencing

*Borrelia burgdorferi* sensu stricto strain N40 was cultured in Barbour-Stoenner-Kelly II (BSK II) medium at +33 °C to a logarithmic phase of growth. Bacteria were washed twice with phosphate-buffered saline (PBS) and counted in a Neubauer cell counting chamber. Bacterial cells were concentrated to 50 x 10^6^ bacteria/ml in PBS. Four female mice (C3H/HeN, aged 4 weeks) were infected by administering 20 µl of the bacterial suspension (10^6^ bacteria/injection) subcutaneously into the hind-leg footpad (Microfine Demi 0.3 ml syringes with a 30-G needle, BD #324826). As a vehicle control, PBS (20 µl) was injected similarly into the other four mice. After four days, mice were sacrificed, and popliteal lymph nodes (draining lymph nodes for the hind leg) were harvested. Lymph nodes from infected mice were pooled into one sample and lymph nodes from control mice were pooled into another sample for lymphocyte isolation. In addition, iliac lymph nodes (further along the chain of draining lymph nodes for the hind leg) were collected and analysed separately with *ospA*-based qPCR to confirm infection. PCR was performed as previously described [12,13].

Single-cell suspension of lymphocytes was produced by mechanically homogenising the pooled popliteal lymph nodes with a custom-made metal cell strainer to detach cells from the stroma. Cells were washed twice with PBS with 2 % fetal calf serum (FCS) buffer and filtered through 77 µm silk. Cells were re-filtered through a 40 µm filter and suspended in RPMI medium with 2 % FCS buffer.

Extracted lymphocytes were processed with the Chromium platform (10X Genomics) for library preparation, and the barcoded scRNA-seq libraries were constructed using 10XGenomics Chromium Next GEM Single Cell 3’, Reagent Kit v3.1. Briefly, reverse transcription was performed to specific single-cell gel beads in the emulsion to produce full-length 10X barcoded cDNA from polyadenylated mRNA. The cDNA was amplified with PCR, followed by enzymatic fragmentation, end repair and A-tailing, adaptor ligation, and sample index PCR. Qualities of cDNA were ensured with Agilent Bioanalyzer 2100, and index PCR was made using Chromium i7 Multiplex Kit. Prepared libraries were sequenced using the Illumina Novaseq6000 Sequencing System (RRID:SCR_016387). Library preparation and sequencing were done in the Finnish Functional Genomics Center at Turku Bioscience.

### 2.2. Bioinformatic analysis

Raw data processing, including demultiplexing, read alignment, and quality control, was performed using the 10X Genomics Cell Ranger package version 3.1.0. Mouse genome, mm10 (GENCODE vM23/Ensembl 98) was used as a reference.

Bioinformatic analysis was performed with R (version 4.3.0 2023-04-21) and RStudio (version 203.03.0). For data analysis, SEURAT – R tool kit for bioinformatic analysis (version 4.0) was used [14].

The control group and *B. burgdorferi*-infected group were processed separately. Low-quality or dying cells, which exhibit extensive mitochondrial gene expression, or low-quality cells with very few genes or none, were identified. Cells with less than 10 % of mitochondrial RNA counts and more than 600 but less than 4000 genes were selected for the analysis.

Datasets from the infection and control groups were normalized and the results were log-transformed. A subset of features that exhibit high cell-to-cell variation were calculated to highlight biological signals. These features that were repeatedly variable across datasets were selected for the integration features. The data were integrated with identified anchors. The integrated data were scaled to ensure that each gene contributes equally to the downstream analysis.

Principal component analysis (PCA) was performed to scale the data to reduce the number of input variables and eventually to make the data easier to visualize. The true dimensionality of the dataset was determined with the JackStraw procedure. Shortly, a subset of the data was rearranged (1 % by default), and PCA was rerun, which constructs a “null distribution” of feature scores and repeats. Fifty dimensions were chosen for the JackStraw procedure. In addition, Elbow plot and dimensionality heatmaps were used to evaluate the lowest PC with biologically relevant information, and PC40 was chosen as a cut-off. Data clustering was constructed as they were iteratively grouped to optimize the standard modularity function. The resolution was set to 0.7 and PC 1:40.

Cell cluster identification was performed with the minimum percentage argument set to 0.4 and the threshold for log2 fold change set to 0.5. Differentially expressed genes between the infection group and the control group were calculated using a Wilcoxon rank sum test. Gene ontology analysis (GO) was made using the Metascape web tool [15].

### 2.3. Re-clustering of T helper cell and B cell subpopulations

The CD4+ T helper and B cell clusters were selected and separately re-clustered with the previously mentioned protocol. Data re-clustering was constructed with the resolution 0.3 and PC 1:20 for B cells and the resolution 0.4 and PC 1:20 for CD4+ T cells. For the cluster identification of CD4+ T cells, the threshold for log2 fold change was set to 0.3, due to the higher similarity of the dataset. Trajectory analysis and ordering cells to pseudotime were performed with Monocle3 [16].

### 2.4. Statistics

The Chi-Square test was used to analyse the significance of cell number differences between the infection and the control group. The Wilcoxon rank-sum test was used to identify the differentially expressed genes and compare the differences in the expression of genes of interest between the infection and the control group with R software. The significance of differentially expressed genes was evaluated by adjusted p-value, based on Bonferroni correction.

### 2.5. Data availability

All raw and processed scRNA-seq data is deposited in the GEO database and are publicly available as of the date of publication [17]. Accession number is GSE248267.

## 3. Results

### 3.1. Immune cell landscape

ScRNA-seq analysis was conducted four days post-infection in a *B. burgdorferi-* infected mouse model, in order to investigate the mechanisms of acute immune response (Fig. 1A). Infection was confirmed with *ospA*-based PCR (data not shown). Altogether 8 840 and 10112 cells were identified in the infection and the control group, respectively.

**Figure 1.**
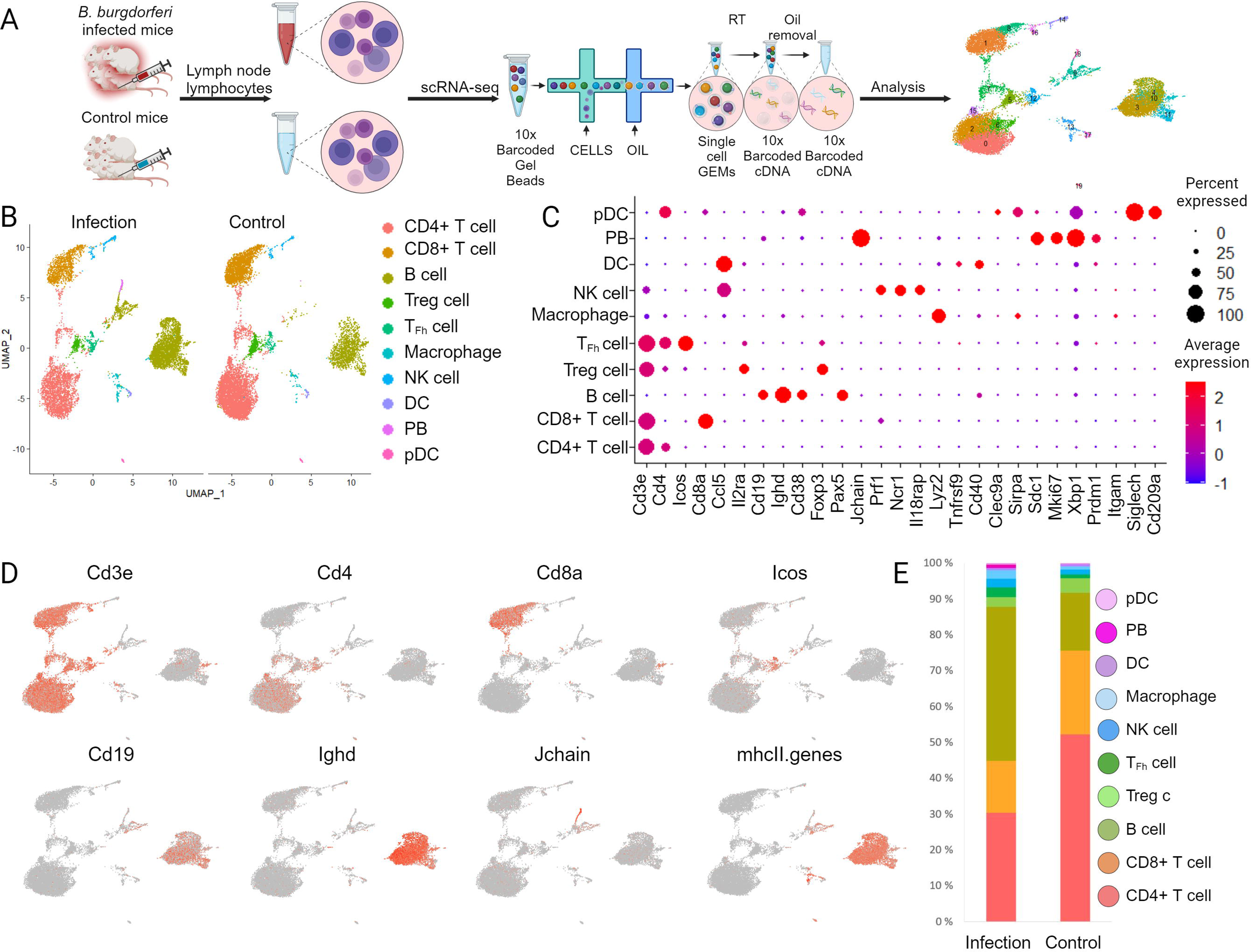
Study workflow and the characterization of the immune cell clusters based on the scRNA-seq data. (A) Experimental outline showing the lymphocyte collection and isolation after four days of infection. Library preparation and sequencing were performed with 10X Genomics® Chromium Single Cell Gene Expression method. (B) UMAP projection of major immune cell clusters from lymph nodes, showing the infection and the control groups respectively. Major cell clusters are shown in different colours. (C) The dot plot representing gene expression profiles of specific marker genes of each major cell lineage. (D) The feature plot showing the expression of selected marker genes across cell clusters. (E) The comparison of relative frequencies of cells of the major lineages in the infection and the control groups. (Abbreviations as follows: Treg: T regulatory, T_Fh:_ T follicular helper, NK: natural killer, DC: dendritic cell, PB: plasmablast, pDC: plasmacytoid dendritic cell)

Clustering of the cells resulted in 20 initial clusters, each representing a different gene expression profile. Based on marker genes, we categorized the cells into ten major cell lineages including CD4+ T cells (*Cd3e, Cd4*), CD8+ T cells (*Cd3e, Cd8a*), T regulatory (Treg) cells (*Cd4, Foxp3, Cd25*), T follicular helper (T_Fh_) cells (*Cd3e*, *Cd4, Icos),* natural killer (NK) cells (*Ccl5, Cd25, Prf1, Ncr1, Il18rap*), B cells (*Cd19, Ighd, Cd38*) PB (*Jchain, Sdc1, Prdm1*), dendritic cells (DC; *Ccl5, Tnfrsf9, Cd40*), plasmacytoid dendritic cells (pDC; *Cd38, Cd11, Siglech, Cd209a*) and macrophages (*Lyz2*)(Fig. 1B-D).

B-cells were significantly more frequent in the infection group, constituting 43.9% (3872 from the total 8840 cells) of the cells, compared to only 16.1% (1633/10112) in the control group (p-value <0.00001) (Fig. 1E). T-cells, particularly CD4+ T-cells, comprised a smaller proportion of the cells in the infection group as compared to the control group (30.4% (2925/8840) vs. 52.2% (5399/10112), p-value <0.00001).

To investigate the extrafollicular response and the previously suggested insufficient T – B cell interaction [5,6], we focused on B cell and CD4+ T cell lineages in the subsequent analysis. Re-clustering of these lineages revealed some misclassified cells (Supplementary Fig. 1 and 2), which were excluded from further analysis.

### 3.2. B cell subset

In the analysis of B cells, in total six different clusters with 3606 cells in the infection group and 1387 cells in the control group were included. (Fig. 2A). These six populations of B cells were identified based on their core markers and changes in the expression levels (↑ indicating upregulation and ↓ indicating downregulation in relation to other clusters) (Supplementary Table 1); naive B cells (*Ighd, Ighm, Ms4a1*), class-switching B cells (*Ighg2b*), proliferating B cells (*Cd19, Mki67*) and PB (*Jchain, Sdc1*) (Fig. 2A-B). We identified two clusters of B cells, which both displayed an activated phenotype, and labelled them as activated I B cells (*Ighd ↓, Ighm ↑, Stat1 ↑*) and activated II B cells (*Ighd ↓, Cd83 ↑*). TACI (*Tnfrsf13b*), a cytokine receptor associated with B cell survival during extrafollicular response, was upregulated in the class switching B cells. The cluster of activated I B cells expressed several interferon-induced genes and, thus, appears to be interferon-activated (Supplementary Table 1). Expression of *Cd83* in the cluster of activated II B cells indicates that the cells are prepared for T cell help, and the gene expression profile suggests that cells are primed for antigen presentation (Fig. 2B) (Supplementary Table 1). The cluster ‘proliferating B cells’ had a more prominent expression of genes related to cell cycle regulation, nucleotide synthesis, and replication. However, the cluster lacked evidence of secretory machinery expansion, or expression of plasma cell transcription factor genes. Nonetheless, the detectable but low expression level of the joining chain of multimeric IgA and IgM (*Jchain)* suggests that these proliferating B cells have initiated the differentiation towards PB and the low expression of *Spi1* and *Irf8* suggests they do not participate in follicular response [18]

**Figure 2.**
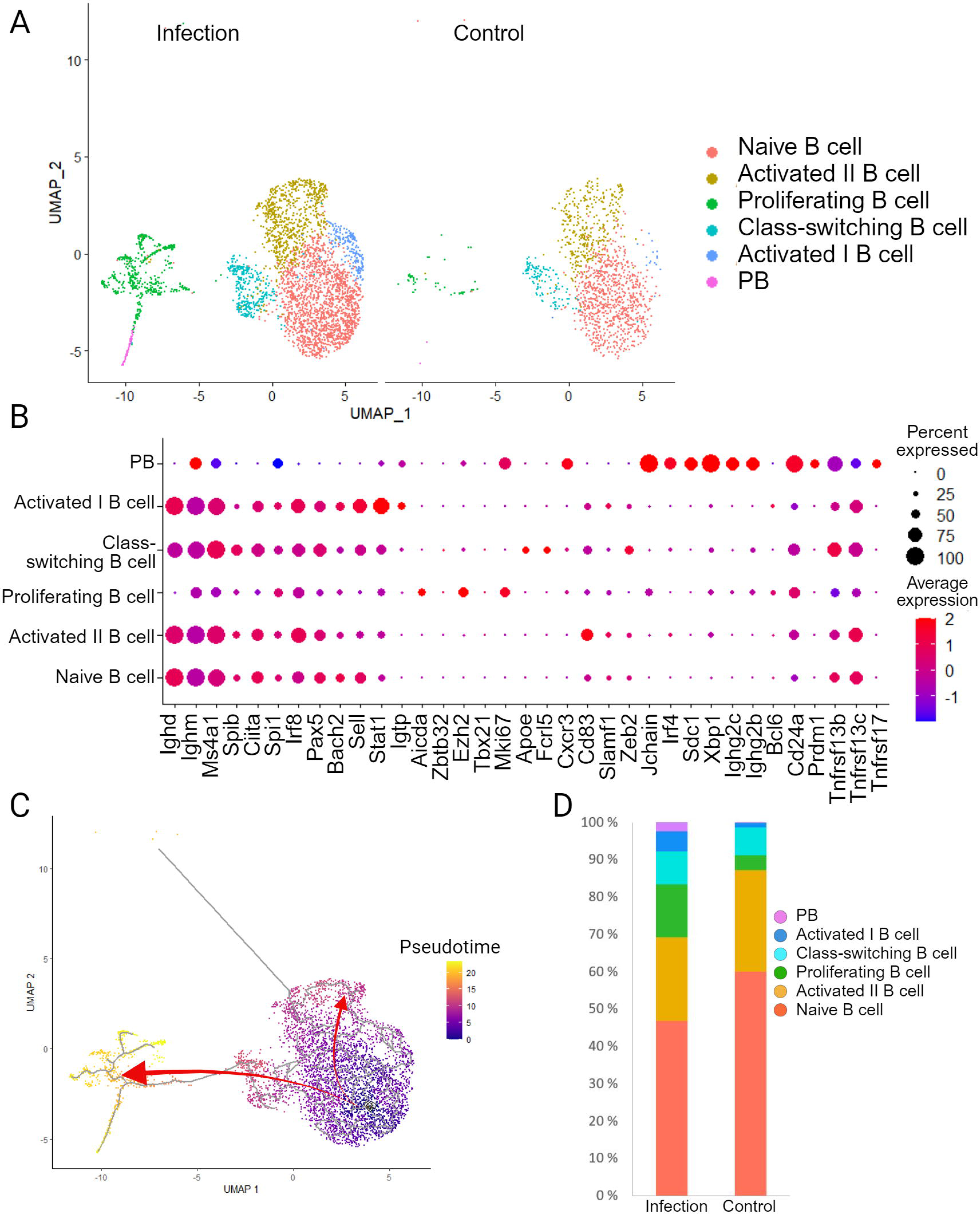
B cell classification and differentially expressed genes between the infection and the control group. (A) UMAP projection of identified B cell clusters, showing the infection and control groups, respectively. Cell clusters are shown in different colours. (B) Dot plot representing the gene expression profile of select marker genes of B cell subtypes. (C) Pseudotime analysis of the B cells suggesting a possible differentiation pathway of extrafollicular immune response (arrow pointing to the left from naive B cells towards the class switching B cells, proliferating B cells and the PB), and of cells that are primed for T cell-dependent response (arrow pointing to up/right from naive B cells towards cluster of activated II B cells). (D) The comparison of relative frequencies of B cell subtypes in the infection and the control groups.

Pseudotime analysis suggests that proliferating B cells and PB represent distinct states compared to other B cell clusters (Fig. 2C). The cluster of activated I B cells was most similar to naive B cells representing similar differentiation stages in pseudotime. Overall, the trajectory analysis and pseudotime plot of the B cell subset suggest two distinct activation pathways, one for the rapid extrafollicular plasma cell differentiation and the other priming for a T cell-dependent response.

The clusters of activated I B cells, proliferating B cells, and PB were significantly enriched in the infection group (p-value <0.00001), whereas the cluster of activated II B cells formed a significantly smaller fraction of the cells (p-value 0.0002) (Fig. 2D). Thus, *B. burgdorferi* infection appears to induce B cell proliferation and PB differentiation via extrafollicular immune response at this time point.

#### 3.2.1. Extrafollicular immune response

To further evaluate the extrafollicular immune response in *B. burgdorferi* infection, we identified 252 differentially expressed genes in the B cell population between the infection and the control groups. The most substantial finding was the significant upregulation of genes associated with plasma cell differentiation (*Jchain*) and immunoglobulin class switching (*Ighg2b, Ighg3*) in the infection group (Fig. 3A-B) (see supplementary Table 2 for significances). Furthermore, we observed that placenta specific 8 (*Plac8*), which is implicated in defence response against bacteria [19,20], and interferon gamma-induced proteasome 20S Subunit Beta 9 (*Psmb9*), important for MHC I antigen presentation [21], were both upregulated in the infection group.

**Figure 3.**
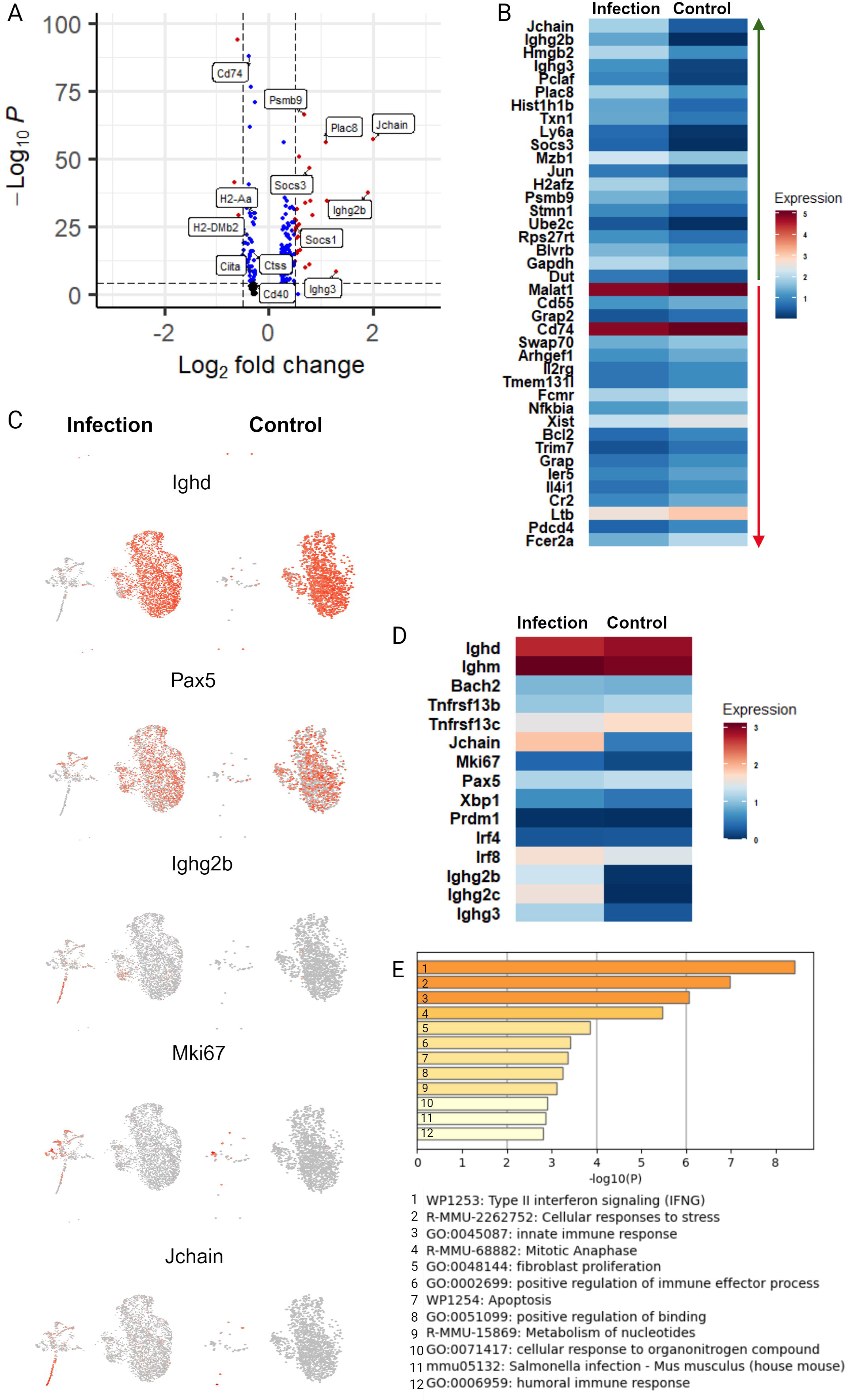
Extrafollicular immune response. (A) A volcano plot showing differentially expressed B cell genes in the infection group compared to the control group. The X-axis represents the fold change between the groups on a log2 scale. The Y-axis shows negative log10 of the p-values. Selected genes are labelled. 252 variables were plotted, black dots represent no significance variation, blue dots represents significance in either Log2FC or p-value, and red dots represent significance for both, Log2FC and p-value. (B) Heatmap of 20 up- and downregulated genes of the infection group compared to the control group. The heatmap includes plasma blast/cell differentiation and immunoglobulin class switching markers that were upregulated in the infection group. (C) Feature plots showing the expression of selected specific differentiation markers from the core markers of B cell clusters, the infection and the control group visualized separately. The feature plots highlight the immunoglobulin class switching, cell proliferation and plasma cell differentiation due to ***B. burgdorferi*** infection. (D) Heatmap of differentiation and isotype switching markers. (E) GO analysis of 40 most upregulated genes of the infection group compared to the control group in all B cells. The graphic is showing the signalling pathways enriched in B cells in ***B. burgdorferi*** infection.

We selected specific differentiation markers from the core markers of B cell clusters and analysed the infection and the control group separately. As expected, the expression of naive B cell marker *Ighd* and the B-lineage specification marker *Pax5*, both expressed at/until the mature stage of the B cell differentiation, were largely similar in infection and control groups (Fig. 3C). This indicates that not all B cells respond to *B. burgdorferi* antigens. However, the increase in infection-induced B cells with the expression of the *Ighg2b*, *Mki67*, and *Jchain* confirms immunoglobulin class switching, cell proliferation, and initiation of plasma cell differentiation. The particularly low expression of *Bcl6*, whose expression is required for initiation and maintenance of germinal center reaction [22], strongly suggests that these populations are not derived from germinal centers (Fig. 2B, Supplementary Fig. 3A). In addition, comparison of differentiation and isotype switching markers show the notable upregulation of extrafollicular immunoglobulin class switching genes in the infection group (*Ighg2b, Ighg2c, Ighg3*) (Fig. 3D).

In conclusion, we observed strong infection-derived immune response in B cells without evidence of germinal center response. Therefore, the gene expression profile of the infection group supports our hypothesis that B cells respond to *B. burgdorferi* infection with an extrafollicular immune response.

#### 3.2.2. SOCS3 and downregulation of MHC II-related genes

To understand how *B. burgdorferi* might promote an extrafollicular response, we focused on the hypothesized regulation of antigen presentation by B cells. Interestingly, our analysis of differentially expressed genes in the infection and the control groups revealed significant infection-derived upregulation of suppressor of cytokine signalling genes *Socs1* and *Socs3* in B cells (Fig. 3A).

We subjected 40 of the most upregulated genes (log2 fold change >0.44) from the B cell subset in the infection group to Gene Ontology (GO) analysis (Fig. 3E, Supplementary Table 3). The analysis revealed significant enrichment of genes associated with Type II Interferon signalling and innate immune response. Interestingly, the observed gene expression pattern paralleled the one reported with *Salmonella* infection (Fig. 3E) [23]. The enrichment summary of the GO analysis revealed negative regulation of tyrosine phosphorylation of STAT protein, negative regulation of receptor signalling pathway via JAK-STAT, and negative regulation of receptor signalling pathway via STAT. Upregulation of three genes, *Socs3*, *Socs1,* and *Irf1,* was involved in the negative regulation of these biological processes, suggesting active inhibition of JAK-STAT pathways of cytokine signalling by Socs proteins.

Alongside the upregulation of *Socs1* and *Socs3*, we saw a significant downregulation of genes that are functionally related to antigen presentation via MHC II (*Ciita, Cd74, Cd40, H2-Aa, H2-DMb2, Ctss*) (Fig. 3A). To investigate whether the downregulation of the MHC II-related genes in the infection group was attributed to PB differentiation, we excluded the proliferating cells and PB from the analysis. Consequently, only subsets with naive and activated B cells (class switching B cells, clusters of activated I and II B cells) were included in this analysis. Despite this refinement, MHC II-related genes continued to exhibit downregulation in the infection group, particularly within the naive B cell cluster (Fig. 4A).

**Figure 4.**
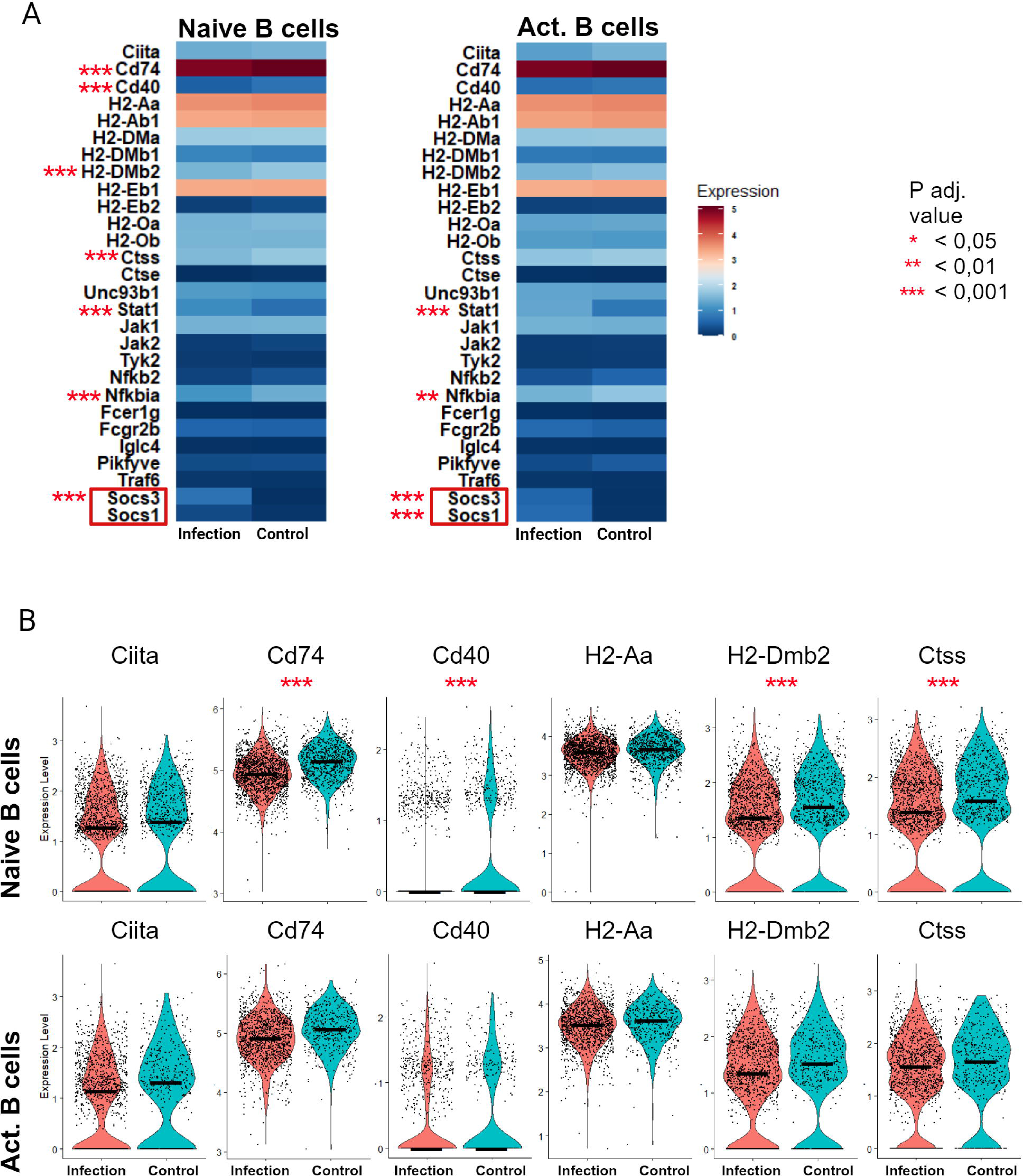
Expression of MHC II -related genes. (A) Heatmap representation of expression levels of MHC II and JAK-STAT pathway-related genes in naive and activated B cells (including class switching B cells, clusters of activated I and II B cells) of the infection and the control group. Significant differences between the groups are pointed out with red asterisks. (B) Violin plot representation of some MHC II related genes in naive B cell and activated B cell populations. Plots with median value visualize the differences in the expression of genes between the infection and the control group. Statistically significant differences between the groups are shown with red asterisks. * = adjusted P value < 0.05; ** = adjusted P value < 0.01; *** = adjusted P value < 0.001

The inhibition of the JAK-STAT signalling pathway is known to reduce MHC II-related gene expression [24]. In agreement with the Socs-mediated downregulation of JAK-STAT pathway activation, violin plots visualize a consistent change in the expression of several genes involved in MHC II-mediated antigen presentation (*Ciita, Cd74, Cd40, H2-Aa, H2-DMb2, Ctss)* between the infection and the control groups, even though all the differences are not statistically significant (Fig. 4B).

Overall, our dataset has no discernible indication of infection-driven upregulation of MHC II-related genes. These results imply that downregulation of the MHC class II transactivator *Ciita* might occur via increased expression of *Socs3* after *B. burgdorferi* infection, which subsequently leads to inhibition of antigen presentation by B cells to T cells.

### 3.3. CD4+ T cell subset

In total, five initial clusters, 2960 in the infection group and 5536 cells in the control group were selected for further analysis of CD4+ T cells. We identified five cell lineages, each representing distinct differentiation stages based on key marker genes; naive CD4+ T cells (*Ccr7, Sell, Klf2, Mif ↓*), Treg cells (*Foxp3*), activated CD4+ T cells (*Cd69, Mif*), T_Fh_ cells (*Icos, Cxcr5, Pdcd1*) and T helper 1 (T_H1_) cells (*Stat1, Ms4a4b*) (Fig. 5 A-B, Supplementary Table 4). Activated CD4+ T cells expressed *Stat6*, suggesting their differentiation into T helper 2 cells. T_Fh_ cells and T_H1_ cells were significantly enriched in the infection group (p-value < 0.00001) (Fig. 5C). The infection group was composed only of 69.8 % (1875 of 2960) of naive CD4+ T cells compared to 87.3 % (4456 of 5536) in the control group (p-value < 0.00001). Treg cells formed relatively the same proportion of cells between the infection and the control group (p-value = 0.0829). We were unable to see whether *B. burgdorferi* infection induces more T_Fh_ cell or T_H1_ cell differentiation, likely due to the early stage of the infection. Trajectory and pseudotime analysis of the CD4+ T cell subset suggests potential differentiation pathways towards activated CD4+ T, Treg, T_Fh_, and T_H1_ cells (Fig. 5 D). Furthermore, T_Fh_ and Treg cells are potentially differentiated earlier compared to T_H1_ and activated CD4+ cells, which is indicated by their differentiation status in pseudotime.

**Figure 5.**
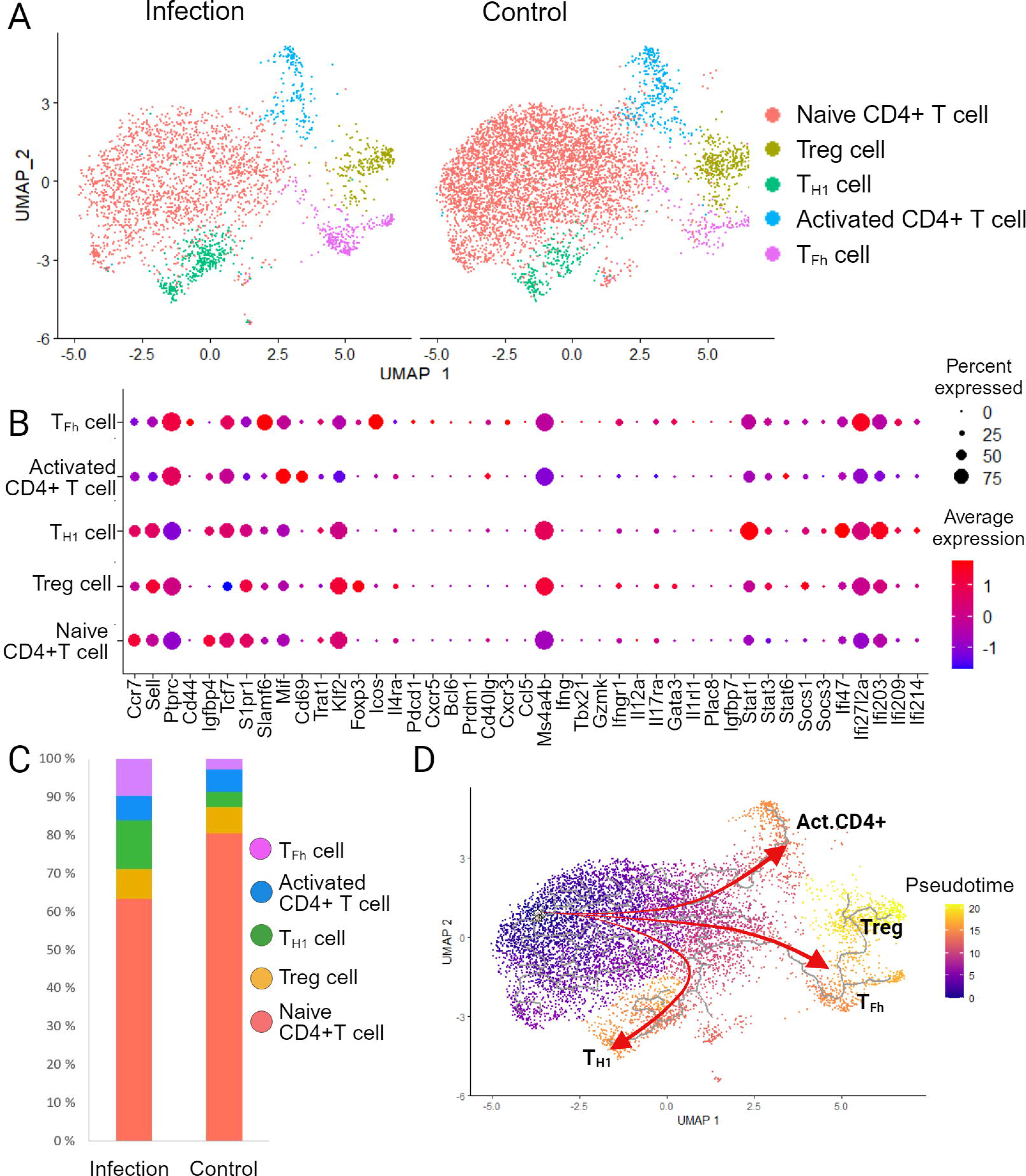
CD4+ T cell subset classification. (A) UMAP projection of identified CD4+ T cell clusters of infection and control groups. Cell clusters are shown in different colours. (B) Dot plot representing the gene expression profile of selected marker genes of CD4+ cell lineages. (C) The comparison of relative frequencies of cells in CD4+ T cell lineages in the infection and the control groups. (D) Pseudotime analysis of CD4+ T cell subset. Red arrows are pointing out possible differentiation pathways of activated CD4+ T, Treg, T_Fh_, and T_H1_ cells from top to bottom, respectively.

Signal transducer and activator of transcription 1 (*Stat1*; p-adj. = 2.8534e-140, Log2FC = 0.8508) and several interferon-induced genes e.g. interferon gamma inducible protein 47 (*Ifi47*; p-adj. = 2.4263e-76 Log2FC = 0.6186) and interferon gamma induced GTPase (*Igtp*; p-adj. = 1.3792e-121, Log2FC = 0.8285), displayed upregulation in the infection group (Fig. 6A). Since *Stat1* is essential for T_H1_ differentiation, its expression may reflect infection-induced T_H1_ cell differentiation. Interestingly, despite the T_H1_ cell differentiation, there was no evidence of *Ifng* (interferon-gamma encoding gene) expression in the infection group (Fig. 5B, Supplementary Fig. 3C). The analysis of significantly changed gene expression of T_Fh_ cells between the control and the infection groups, suggested that some of the T_Fh_ cells were prepared for germinal center responses in the infection group (Fig. 6B). *Il21*, the gene encoding a cytokine that is critical for germinal center formation (p-adj. = 3.4748e-04, Log2FC = 0.5447), and *Cxcr5*, the gene encoding chemokine receptor 5 that localizes T_Fh_ cells to developing germinal centers (p-adj. = 2.3174e-07, Log2FC = 0.5496), were upregulated by T_Fh_ cells in the infection group. While the differences were statistically significant, only a small pool of T_Fh_ cells in the infection group showed expression of *Il21* and *Cxcr5*.

**Figure 6.**
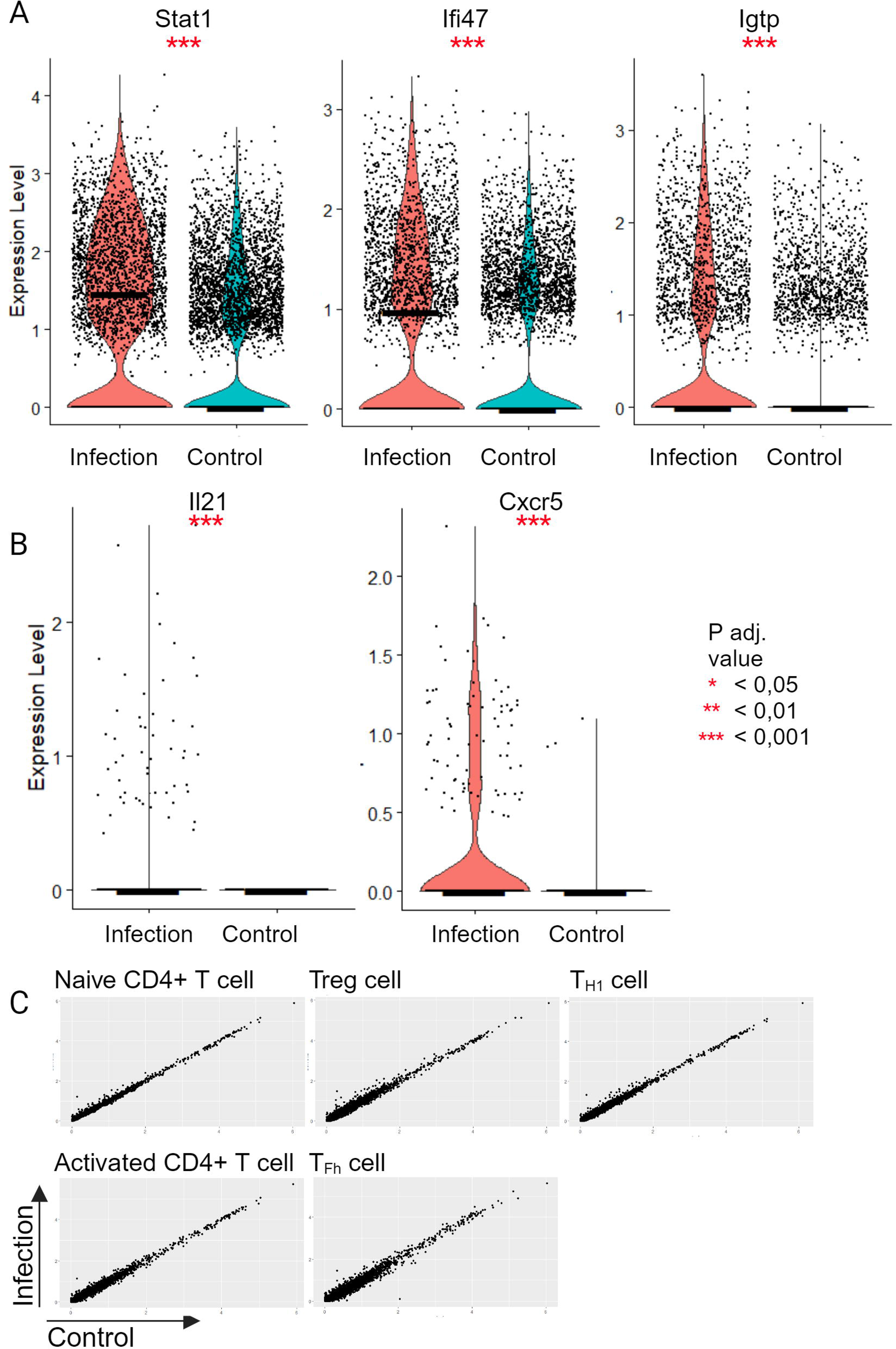
Differentially expressed genes in CD4+ T cells between the infection and the control group. (A) Violin plot representation with medians of ***Stat1*** and some interferon-induced genes (***Ifi47, Igtp***), which were significantly upregulated in the infection group. Significances are pointed out with red asterisks. (B) Violin plot representation with expression of ***Il21*** and ***Cxcr5*** in T_Fh_ cell subset of the infection group compared to the control group. (C) Scatter plots representing the average gene expression of CD4+ T cell clusters in the infection and the control groups, highlighting the similarity of the datasets despite the infection. X-axis represents the average gene expression of the control group and the Y-axis represents the average gene expression of the infection group. * = adjusted P value < 0.05; ** = adjusted P value < 0.01; *** = adjusted P value < 0.001

In conclusion, CD4+ T cells reflected broadly similar transcriptomic profiles in the infection and in the control group (Fig. 6C). Only a few cells displayed detectable expression of cell proliferation marker *Mki67*, indicating that the CD4+ T cell population was not significantly activated (Supplementary Fig. 3D). However, the increases in T_Fh_ and T_H1_ cells suggest that there is some activation due to *B. burgdorferi* infection.

## 4. Discussion

This study describes scRNA-seq profiling of lymph node lymphocytes in *B. burgdorferi* infection in mice. We focused on the early immune response, which primarily involves extrafollicular B cell responses, and found infection-induced upregulation of suppressor of cytokine signalling genes *Socs1* and *Socs3* in B cells. Our results suggest a model where *Socs3*-mediated regulation restricts T – B cell interaction in early B*. burgdorferi* infection.

In support of our hypothesis of a limited B and T cell interaction, earlier studies of mice infected with *B. burgdorferi* have demonstrated a failure in the development of protective immunity after vaccination with influenza antigens, underscoring the inhibitory effect of *B. burgdorferi* on the development of long-term humoral immunity [6]. The exact molecular mechanisms behind the phenomenon have remained obscure, even though it has been studied broadly for example with flow cytometry and histology [5,9,22,25].

In recent years, the heterogeneity of gene expression within cell populations has become increasingly evident [26,27]. ScRNA-seq approach has been successfully applied to analyse for example the heterogeneity of tumours and cellular developmental pathways with high resolution. Studies of single-cell transcriptomic profiles in infection immunity have remained relatively limited but have underscored, e.g., a key role for B cells in response to *B. burgdorferi* in the erythema migrans lesions, and revealed transcriptional changes following *B. burgdorferi* infection in murine ankle joints [28,29].

Using single-cell RNA sequencing, we found that B cells were highly abundant in mouse lymph nodes after infection, while T cells represented a smaller proportion. A rapid enlargement of the regional inguinal lymph nodes was previously observed starting at day four in *B. burgdorferi*-infected C57BL/6 mice, where the increased lymph node cellularity was shown to be due to a massive expansion of B cells [9]. The accumulation of B cells may disturb germinal center formation, and the lack of functional germinal centers or failure to maintain them results in the inability to generate long-lived plasma cells and efficient B cell memory [6,11,22]. Indeed, Hastey et al. found a clear demarcation of T and B cell zones only up to day four of the *B. burgdorferi* infection, and the increase in B cell frequency was accompanied by a corresponding decrease in CD4+ T cell frequency [25]. In line with our results, CD4+ T cells have been shown to differentiate into T follicular helper phenotypic cells following *B. burgdorferi* infection, suggesting that there is not a total defect in T cell responses, but rather T-dependent responses are suppressed, which could contribute to the mainly extrafollicular immune response [6,11].

We sought insight into the reasons behind the mainly extrafollicular *B. burgdorferi*-specific B cell responses, focusing on the early time point of the infection [22]. In this model, we saw a clear activation and accumulation of B cells in *B. burgdorferi* infection consistent with lymphadenopathy associated with *B. burgdorferi* infection in mice [9]. Accumulation was also visible in the naive B cell cluster. Our results indicate that within just four days, *B. burgdorferi* infection triggers a notable B cell proliferation, immunoglobulin class switching to IgG3 and IgG2b isotypes, and induces plasma cell differentiation, all hallmarks of extrafollicular immune response [10]. The expression of TACI (*Tnfrsf13b*) further suggests that immunoglobulin class switching is induced during the extrafollicular immune response [30]. Interestingly, the most upregulated genes of B cells of the infection group have similarity to the reported gene expression patterns in mice during *Salmonella* infection, which is also known to trigger an extrafollicular response [23]. Importantly, there was no evidence of high *Bcl6* expression in any of the B cell clusters, indicating the lack of germinal center response. These findings are in line with previously described predominantly extrafollicular responses to *B. burgdorferi* [4,9].

We found that both T_H1_ and T_Fh_ cell populations were more abundant in the infection group compared to the control group and the upregulation of *Il21* and *Cxcr5* in T_Fh_ cells suggests that some of the cells were instructed for germinal center reactions. Strong T_H1_ differentiation with the lack of T_Fh_ cells would have the potential to explain why the immune response during *B. burgdorferi* infection fails to maintain T-dependent germinal center responses. However, the early stage of the infection does not allow for the determination of whether *B. burgdorferi* infection primarily drives T_H1_ or T_Fh_ cell differentiation. Previous findings suggest that *B. burgdorferi* infection is T_H1_ dominant [31,32]. Four days after infection, we could not see any induction of *Ifng* (encoding interferon-gamma) expression, despite the seemingly initiated differentiation into T_H1_ cells. Our data suggest that interferon-gamma is rather secreted by other cells than the T_H1_ cell population. However, neither natural killer cells, macrophages nor dendritic cells of our dataset showed expression of *Ifng* (data not shown). Hammond et al. 2023 have reported similar results recently in a later time point [11]. Their data suggest that *B. burgdorferi* infection does not induce interferon-gamma producing CD4+ cells in lymph nodes.

Earlier, *B. burgdorferi* infection has been hypothesized to induce restricted T cell responses, possibly through increased expression of *Socs3* in dendritic cells, macrophages, and monocytes leading to reduced antigen presentation to T cells [33,34]. In macrophages, *B. burgdorferi* spirochetes or lipidated outer surface protein A (*L-OspA*) increases *Socs1* and *Socs3* mRNA and protein expression [34]. Such an effect has not been described in B cells until now. Our data indicate that a Socs-mediated downregulation of the antigen presentation pathway may also be active in B cells.

In the context of many bacterial infections, *Ciita* expression is induced in B cells. CIITA is a major transcription factor promoting the expression of the MHC II molecules during antigen presentation to T helper cells [35]. *Ciita* transcription, in turn, is directly promoted by STAT1 [36]. Contrary to our expectations, we did not observe infection-driven upregulation of MHC II-related genes. Instead, many of the genes, including *Ciita*, were consistently downregulated, although not all the differences reached statistical significance. One plausible interpretation arising from this analysis is that the observed negative regulation of receptor signalling pathways via STAT/JAK-STAT in *B. burgdorferi* infection results from *Socs1* and *Socs3* upregulation. These inhibitors of JAK-STAT signalling could cause the downregulation of genes of MHC II antigen presentation, and of other genes regulating B-T cell interactions and thus preventing B cells from receiving adequate T cell help. These findings may provide a partial explanation for why *B. burgdorferi* directs the immune response toward the extrafollicular B cell pathway. Our results suggest that the impaired germinal center responses in *B. burgdorferi* infection are driven by a strong extrafollicular response and/or restricted B - T cell communication.

The immune evasion strategies of *B. burgdorferi* are diverse [37]. During a tick bite, tick saliva inhibits lymphocyte proliferation and antibody production by B cells [38]. Infection via needle inoculation is commonly used in *B. burgdorferi* infection studies and may preclude some immune evasion mechanisms of natural tick-borne infection. We used young C3H/HeN mice since they show clear signs of inflammation and develop arthritis in *B. burgdorferi* infection [39], however, the complete maturation of their immune system was not controlled. Overall, our results are parallel with typical immune response patterns described in tick-borne *B. burgdorferi* infections [9]. In addition, our results allow the identification of a new potential suppressive mechanism of T lymphocyte activation in *B. burgdorferi* infection.

Taken together, our results highlight the role of B cells in the immune response to *B. burgdorferi* infection. The gene expression patterns observed at the early time-point of the infection suggest a predominantly extrafollicular immune response characterized by the secretion of IgG2b and IgG3 by short-lived plasma cells. In the future, analysis of T and B cell interactions during *B. burgdorferi* infection at later time points and with control infections would provide a more comprehensive view of the balance between extrafollicular and germinal center responses.

## Ethics approval

All animal experiments were approved by the Ethical Committee for Animal Experimentation in Finland. Experiments were performed according to the Finnish Act on Animal Experimentation (497/2013) in compliance with the 3R-principle with the appropriate animal license (ESAVI/6265/04.10.07/2017).

## Acknowledgements

This work was supported by the Special Governmental Fund for University Hospitals. We thank Finnish Functional Genomics Centre supported by University of Turku, Åbo Akademi University, and Biocenter Finland and Single Cell Omics Facility of Turku Bioscience. Figures of this article were created with Biorender.com

