## Supplementary Figure1 for "Single-cell transcriptome analysis of the early immune response in the lymph nodes of *Borrelia burgdorferi*-infected mice"

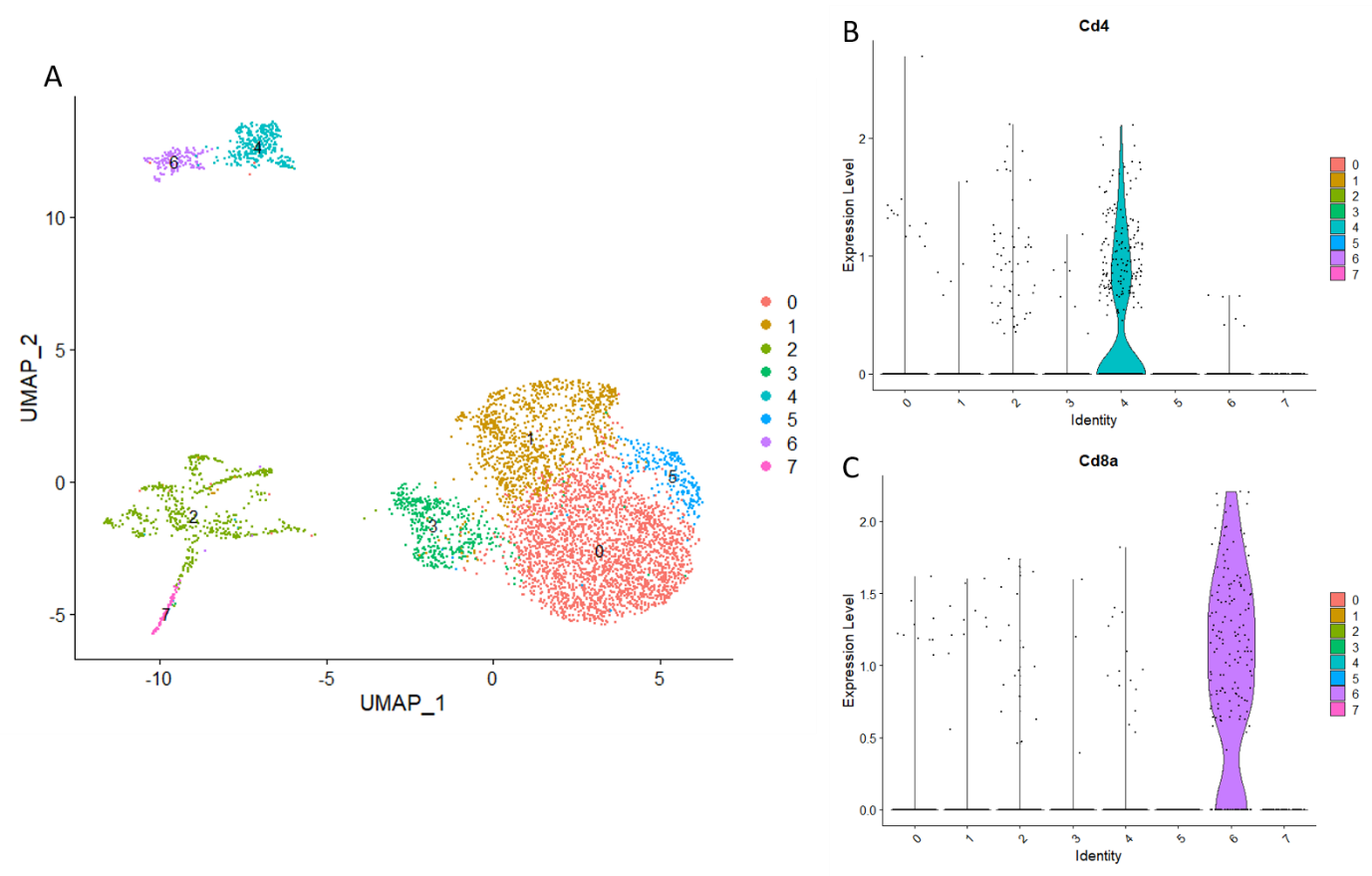

Supplementary Figure 1. Re-clustering the B cell subpopulation.

(A) Re-clustering of B cells resulted in eight distinct clusters. (B) Cluster 4 was identified as Cd4+ cell contamination. (C) Cluster 6 was identified as Cd8a+ cell contamination.
