## Supplementary Figure2 for "Single-cell transcriptome analysis of the early immune response in the lymph nodes of *Borrelia burgdorferi*-infected mice"

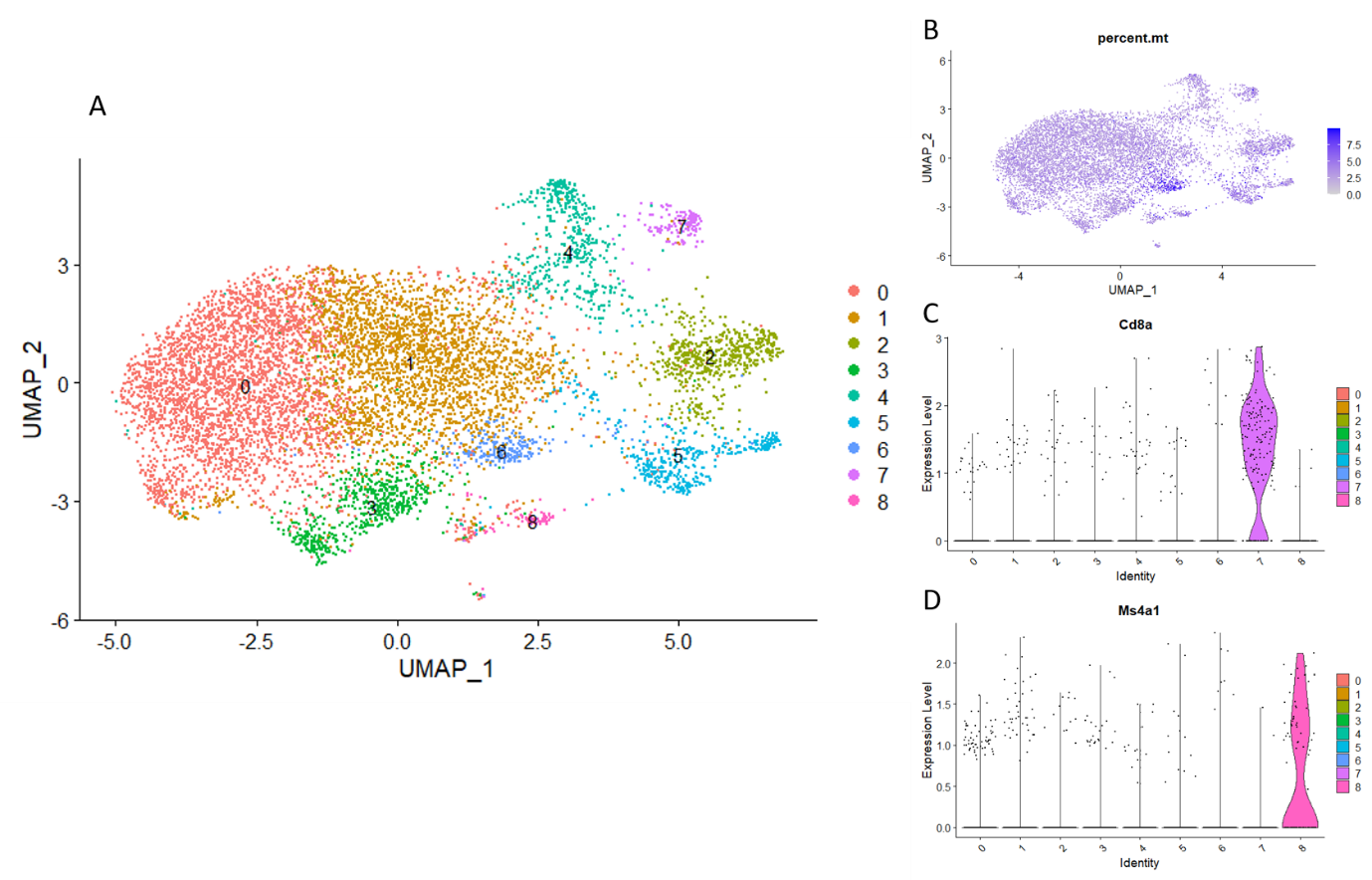

Supplementary Figure 2. Re-clustering the CD4+ cell subpopulation.

(A) Re-clustering of CD4+ cells resulted in nine distinct clusters. (B) Cluster 6 expressed mitochondrial DNA and was considered to comprise of dying cells. (C) Cluster 7 was identified as *Cd8a*+ cell contamination. (D) Cluster 8 was identified as B cell contamination (*Ms4a1*)
