## Supplementary Figure3 for "Single-cell transcriptome analysis of the early immune response in the lymph nodes of *Borrelia burgdorferi*-infected mice"

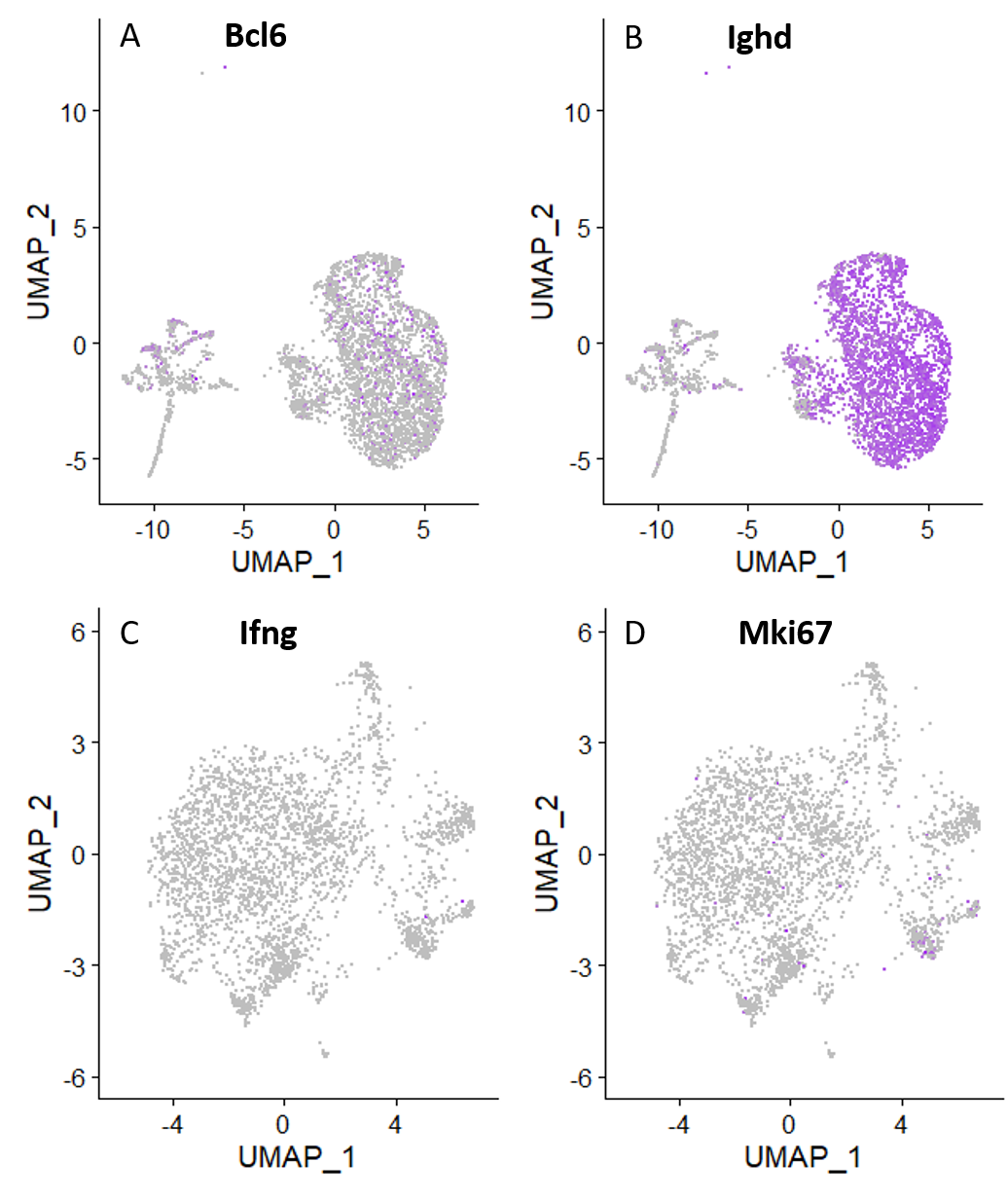

Supplementary Figure 3. Expression of selected genes in the infection group.

(A) Low expression of *Bcl6* and (B) high expression of *Ighd* in the B cell subset of the infection group. (C) Absence of the *Ifng* expression and (D) low expression of *Mki67* in the CD4+ T cell subset of the infection group.
